# Thorax temperature and niche characteristics as predictors of abundance of Amazonian Odonata

**DOI:** 10.1101/2024.09.14.613059

**Authors:** Lenize Batista Calvão, Ana Paula J. Faria, Carina Kaory Sasahara de Paiva, José Max Barbosa Oliveira-Junior, Javier Muzón, Alex Córdoba-Aguillar, Leandro Juen

## Abstract

Environmental architecture and body temperature drive the distribution of ectothermic species, especially those with specific ecophysiological requirements or narrow ecological niches. In this study, we evaluated the connection between thorax temperature and niche specialization concerning the abundance and species contribution to the beta diversity of adult Odonata in Amazonian streams, employing the Species Contribution to Beta Diversity (SCBD). Our hypotheses were (i) Odonata species’ thorax temperature is positively correlated with both morphology (thorax width) and air temperature, and (ii) the thorax temperature of the Odonata assemblage serves as a more influential predictor than niche specialization in determining species abundance and SCBD. We sampled 46 streams in an anthropized landscape in the Northeastern and Southeastern regions of Pará state, Brazil. Notably, niche breadth emerged as the variable influencing the abundance and SCBD of the Odonata assemblage. Niche position is a predictor for Odonata SCBD and not suborders, and predictor for abundance, except for Anisoptera. Both suborders exhibited a negative relationship between abundance and thoracic temperature. In summary, our results underscore the necessity of considering both niche and ecophysiological predictors to comprehensively assess the Odonata assemblage in Amazonian streams. This holistic approach has implications for conservation efforts and bioassessment practices, offering valuable insights into the collective response of Odonata as a group.

## Introduction

One aim in ecology is to understand how species assemblages distribute as a function of each species’ requirements. In this regard, ecological niche breadth and position are key predictors for the local abundance [1–4]. The specific link that underlies species abundance and such predictors is explained according to two models. The hypothesis that links niche breadth/tolerance, predicts that populations that can remain viable in a wide range of environmental conditions have greater abundance. In this context, they can be characterized as generalists (occur in a wide range of environmental conditions) and specialists (occur in a more restricted range). Conversely, the hypothesis for the niche position predicts that non-marginal species, those that occur in a larger availability of habitats, tend to be more abundant(11).

Generalist species, endowed with greater niche breadth, can occur in a wider range of environmental conditions, leading to a smaller contribution of species for Beta diversity (SCBD) than with the contribution of specialist species [10]. Accordingly, niche position can influence SCBD, as species in marginal habitats use more specific environmental conditions than non-marginal species [10]. Simultaneously, the relationship of functional traits (e.g., thermoregulatory ability governed by body size) with SCBD may occur if traits confer adaptations in the species, influencing their abundance and/or occupancy at the site [2, 4].

Odonata are aquatic insects whose thermal performance determines their distribution both at the micro-[12] and macro-scale [13]. In this context, the body size and behavior of adult odonates are associated with their thermoregulation strategies, categorizing species as thermal conformers, heliothermic or endothermic [12]. These thermoregulatory strategies are correlated with body size, as heat exchange with the environment occurs based on the surface/volume ratio [14]. Consequently, small percher species (e.g., most zygopterans) generally exhibit thermal conformers or heliothermic, relying on air temperature to start their activities [12]. These species can regulate heat loss by adjusting their body posture in response to light [15–16] and by selecting habitats that support their thermal requirements [14, 17]. Conversely, larger species of Odonata can inhabit open areas with reduced canopy cover [12] as they are heliothermic [18], benefiting from direct sunlight exposure. Finally, larger species (e.g., mostly Anisoptera) are predominantly endothermic, enabling them to overcome the limitations faced by thermal conformers, as they can internally control heat loss or production through their thorax muscles [14].

Ecophysiological requirements have frequently been crucial predictors for odonate abundance or distribution in tropical streams [18, 19, 20, 21]. This correlation is primarily supported by the impact of changes in vegetation cover and air temperature, both acting as environmental filters influencing odonate community composition [21]. For example, the absence of vegetation and reduced shade favor most anisopteran species due to their reliance on increased light input into the stream [12, 18, 20, 21, 22]. Conversely, these environmental conditions are unsuitable for most Zygoptera or smaller species, as they typically prefer habitats with a more consistent temperature and may not be able to thermoregulate effectively in altered streams [20, 21, 22].

In the present study, we aimed to evaluate the importance of thoracic temperature and niche specialization on the abundance and contribution of each species to the SCBD of adult Odonata in Amazonian streams. For this, we had the following predictions: (i) thoracic temperature of Odonata species differs between suborders and correlates positively with morphology (thorax width) and air temperature. Thus, larger species (Anisoptera) may exhibit heliothermic behaviors, leading to thoracic temperature higher than air temperature. Conversely, smaller species (Zygoptera) tend to thermoregulate in response to air temperature, resulting in thorax temperature similar to the ambient environment; (ii) the thoracic temperature of the Odonata assemblage, as well as its suborders (Zygoptera and Anisoptera), serves as a more important predictor than niche characteristics in predicting the abundance and SCBD of the species given that ecophysiological traits play a crucial role in Odonata habitat selection (Fig 1).

**Fig 1.**
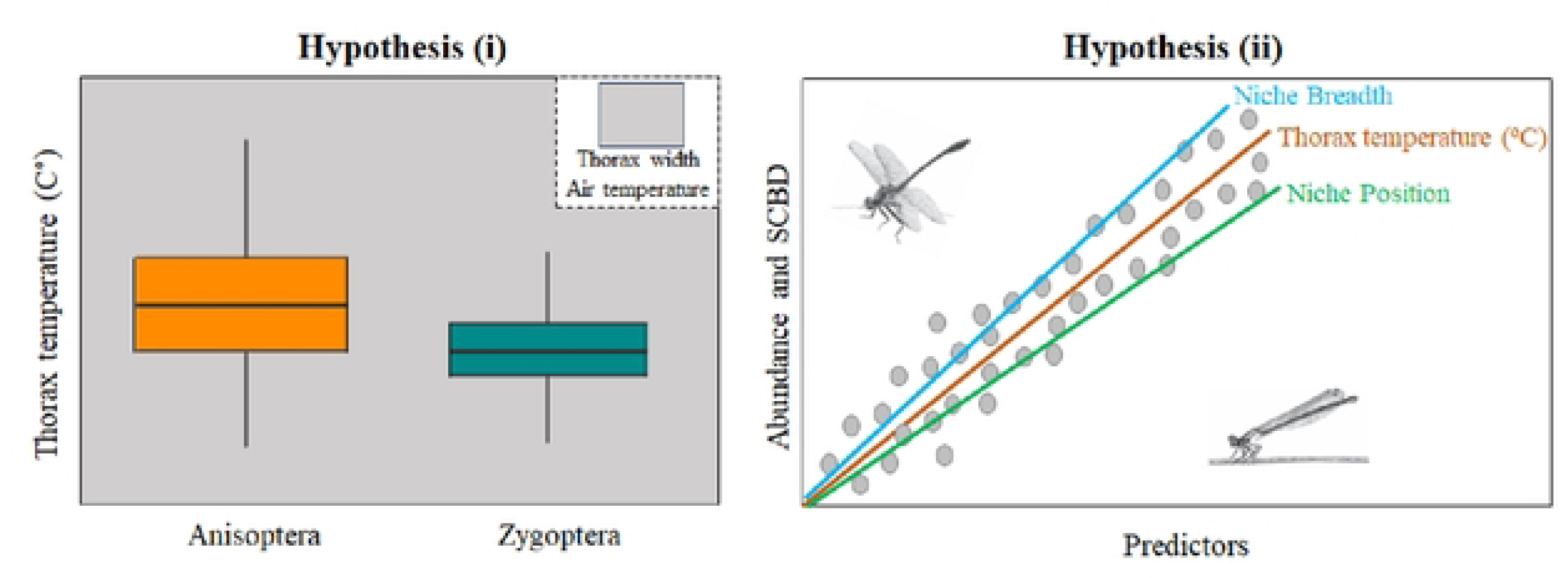
Schematic drawing of hypotheses and images of Odonata were adapted from Stehr, F.W and Tennessen, K.J.

## Material and methods

### Study area

The study was conducted in 46 streams across four municipalities in the Northeastern region (Tomé-açu, Ipixuna do Pará, Concórdia do Pará and Acará) and three municipalities in the Southeastern region (Paragominas, Canaã dos Carajás and Parauapebas) of Pará state, Brazil, covering basins ranging from the Tocantins-Araguaia River and Capim River (Fig 2, S1 Table). Northeastern Pará is characterized, according to the Köppen classification, by a tropical rainforest climate (Af) and tropical monsoon climate (Am) [23], with temperature ranging from 22°C to 34°C (minimum: 22°C to 23°C; maximum: 30°C to 34°C). Southeastern Pará has a Savanna climate, classified as a tropical climate with a dry season (Aw) [23, 24, 25]. The municipalities of Canaã dos Carajás and Parauapebas, located in Serra dos Carajás area, are notable for their landscape, mainly due to their elevation, which ranges between 500 and 700 m a.s.l. [24]. The average monthly temperature in this area varies from approximately 25°C to 26°C, with an annual rainfall of 2.033 millimeters [24].

**Fig 2.**
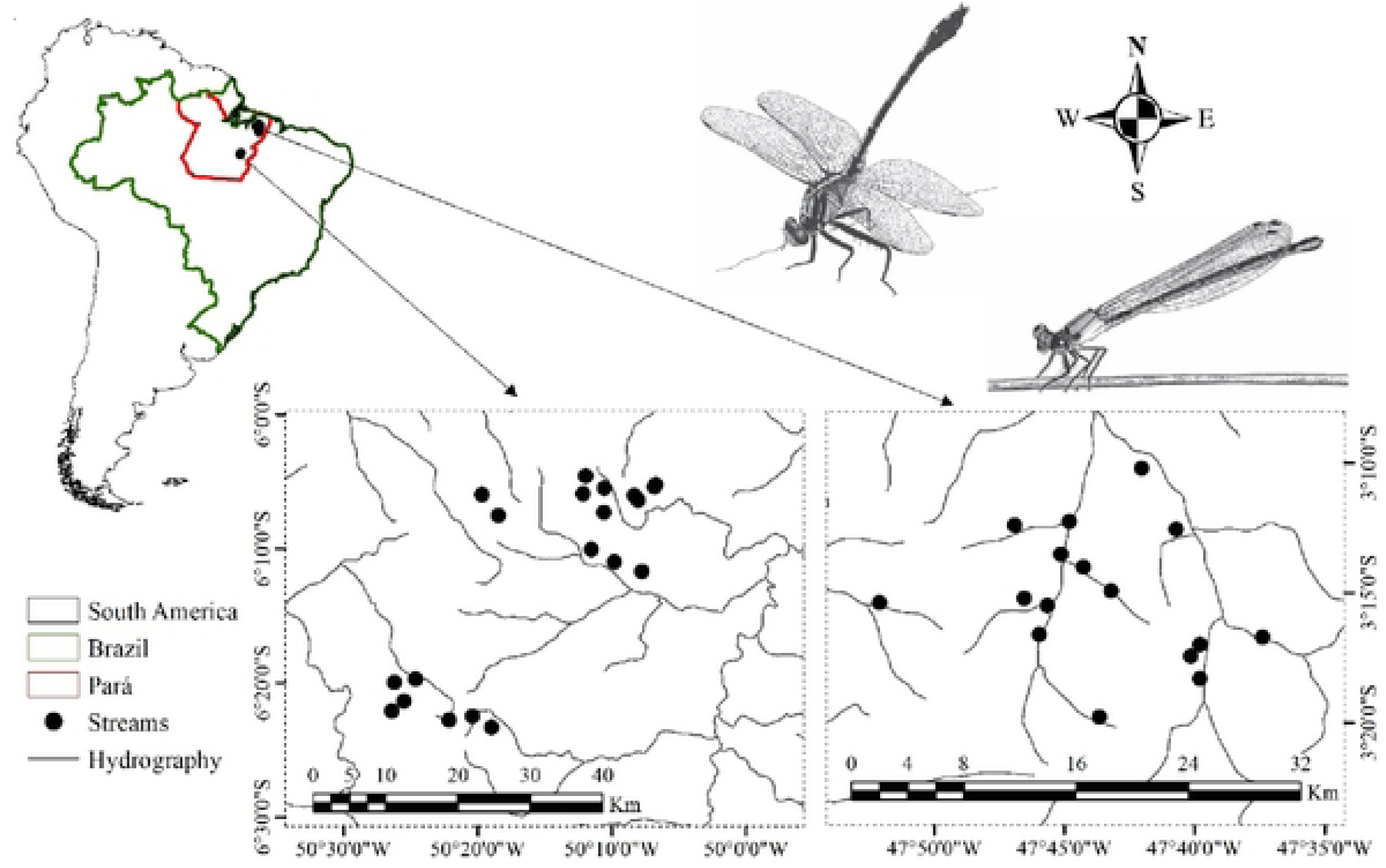
Study area showing the 46 streams distributed in the Northeastern and Southeastern of Pará, Brazil. Images of Odonata were adapted from Stehr, F.W and Tennessen, K.J.

### Insect sampling

The 46 sampled streams ranged from 1st to 3rd order, with an average width of 2.8 meters and a depth of 29.9 centimeters, according to Strahler [26] classification. Sampling periods were performed between July 2017 and October 2018, consistently during the low precipitation period. Adult odonates were collected only once stream, in 20 segments spaced 5m apart, distributed along a continuous 150-meter longitudinal stretch in each stream [further details in 20, 27, 28]. Specimens were collected on stream banks for one hour, always between 11:00 and 12:00 hrs (the peak activity period for adult odonates), at temperatures above 19°C, using a 40cm diameter and 65cm in length entomological net [21, 29]. Standardization of sampling effort and climatic conditions was necessary due to the organisms’ thermoregulatory ability based on solar incidence [30], ensuring the presence of all ecophysiological groups [21].

The collected specimens were placed in tracing paper envelopes and subsequently fixed in acetone P.A. (Propanone) for 12 hours for Zygoptera and 48-72 hours for Anisoptera. Species identification was conducted to the species level using specialized taxonomic keys [31–35]. The biological material was deposited in the Collection of Aquatic Insects at the Laboratory of Ecology and Conservation, Federal University of Pará (UFPA), University Campus of Belém, Pará, Brazil [36].

### Measurements of physiological traits

The thoracic temperature of males was captured using an entomological net and quickly measured within a maximum of 10 seconds to avoid physical damage and alteration in body temperature. We used only male individuals because species identification is possible in this sex. Holding the wings, we measured the body temperature (°C) of each individual using a Testo-805 infrared thermometer (accuracy ± 1°C [-2 to +40°C], resolution 0.1°C [-9.9 to +199.9°C], and reaction time <1s). The recording was performed by pointing the thermometer beam at the center of the right side of the thorax, at a distance of five centimeters.

Thorax width was measured using a digital caliper (accuracy = 0.02 mm) for males of each species. For species with multiple individuals, we used the averages, while for those with only one individual, we recorded the absolute value.

### Environmental variables in streams

For each stream, four environmental variables were measured: depth (cm), channel width (m), Habitat Integrity Index (HII) and air temperature (in °C). Depth was measured at three points of the stream: left and right of the bank, and central region. The average of these measurements was used as a measure of depth in each sampling unit. The HII was composed by 12 items that describe the degree of habitat integrity in the stream: land use pattern beyond the riparian zone; width of riparian forest; completeness of riparian forest, vegetation of riparian zone within 10 m of channel,; retention devices and sediment in the channel; river bank structure; bank undercutting, stream bottom; distribution of riffles and pools; characteristics of aquatic vegetation and detritus [37]. Each item presented four to six alternatives corresponding to the observed condition related to habitat integrity. We transformed each item value to produce the HII, which ranges from 0 (altered stream) to 1 (preserved stream), according to the habitat integrity conditions found in each stream [37]. Note that this HII has been widely used in studies that have assessed the environmental conditions of streams and their relationship with aquatic insect diversity [38–40]. Air temperature was measured using a Testo-805 infrared thermometer. After measuring the thorax temperature of the specimens, the thermometer was positioned in the microhabitat where each individual was collected.

### Data analysis

To calculate niche breadth and position (i.e., predictor variables) of the species, we used the Outlying Mean Index (OMI [11]) which is based on the following environmental variables: depth, channel width, and HII. The OMI analysis calculates the distance between the average environmental conditions used by the species (centroid) and the average environmental conditions of the sampled sites (hyperspace) [11,41]. Species abundance data were logarithmized and environmental variables standardized. The marginality significance (OMI) of the species was evaluated using the Monte-Carlo permutation test with 1000 permutations [11]. Thus, we obtained the breadth (environmental tolerance) and position (marginality) of the ecological niche for each species. We carried out PCoA for Odonata composition visualization using Hellinger and Bray-Curtis.

The ecological uniqueness of species (SCBD) was measured from the total variation of the assemblage (Total Beta Diversity - BD_Total_) between streams. For this analysis, we submitted the Odonata composition matrix to the Hellinger transformation, which was used to measure the Sum of Total Squares (SS_Total_). Then, we divided the SS_Total_ by the number of sampled streams (n-1) and obtained the BD_Total_ of the assemblage that was partitioned into ecological uniqueness of species (SCBD) [9]. For more details on the calculation of BD_Total_ and SCBD, see [9]. Finally, species with higher SCBD values had a higher relative contribution to beta diversity [9].

We performed the analysis of the relationship between the variables from the generalized linear mixed models (GLMMs) and species as random factors. Thus, to assess (i) the difference in thoracic temperature of Anisoptera and Zygoptera, the relationship between the interaction of air temperature and thorax width on Odonata thoracic temperature, and the relationship between the difference of air and thoracic temperature (Tair-Tth [°C]) and thorax width for Anisoptera and Zygoptera, with Gaussian distribution. The analyzed models contained each individual as a sample unit.

To evaluate (ii) the effect of thoracic temperature, niche breadth and position (predictor variables standardized) on the abundance (response variable) of Odonata species, we carried out a GLMM and suborder as a random factor, using the negative binomial distribution due to data overdispersion [42]. For this model we used the Log linkage function. The analyzed models contained species as a sample unit. For Suborders, we carried out a GLM (Negative Binomial). We performed the visual validation of the models using the simulated envelope [43]. To assess the effect of thoracic temperature, niche breadth and position (predictor variables standardized) on Odonata and suborders SCBD (response variable), we used a Beta Regression [44]. This analysis is more suitable for the response variable (SCBD) distributed between values from 0 to 1 [44]. The binding function used in this model was logit.

All analyses were performed using the R software [45] using MASS [46], hnp [43], betareg [44], and vegan packages [47].

## Results

### Odonata assemblage

859 specimens were collected, belonging to 56 species: 22 anisopterans and 34 zygopterans (S2 Table).

### Thoracic temperature and size, and their relation with ambient temperature

Anisopterans showed greater temperature and thorax width than zygopterans. On average, Anisoptera species have 5°C more than Zygoptera. Air temperature was higher 2°C in the places where Anisoptera species were sampled (Table 1, Table 2 and Fig 3). Both thoracic width and air temperature affect thoracic temperature, and (Table 2).

**Fig 3.**
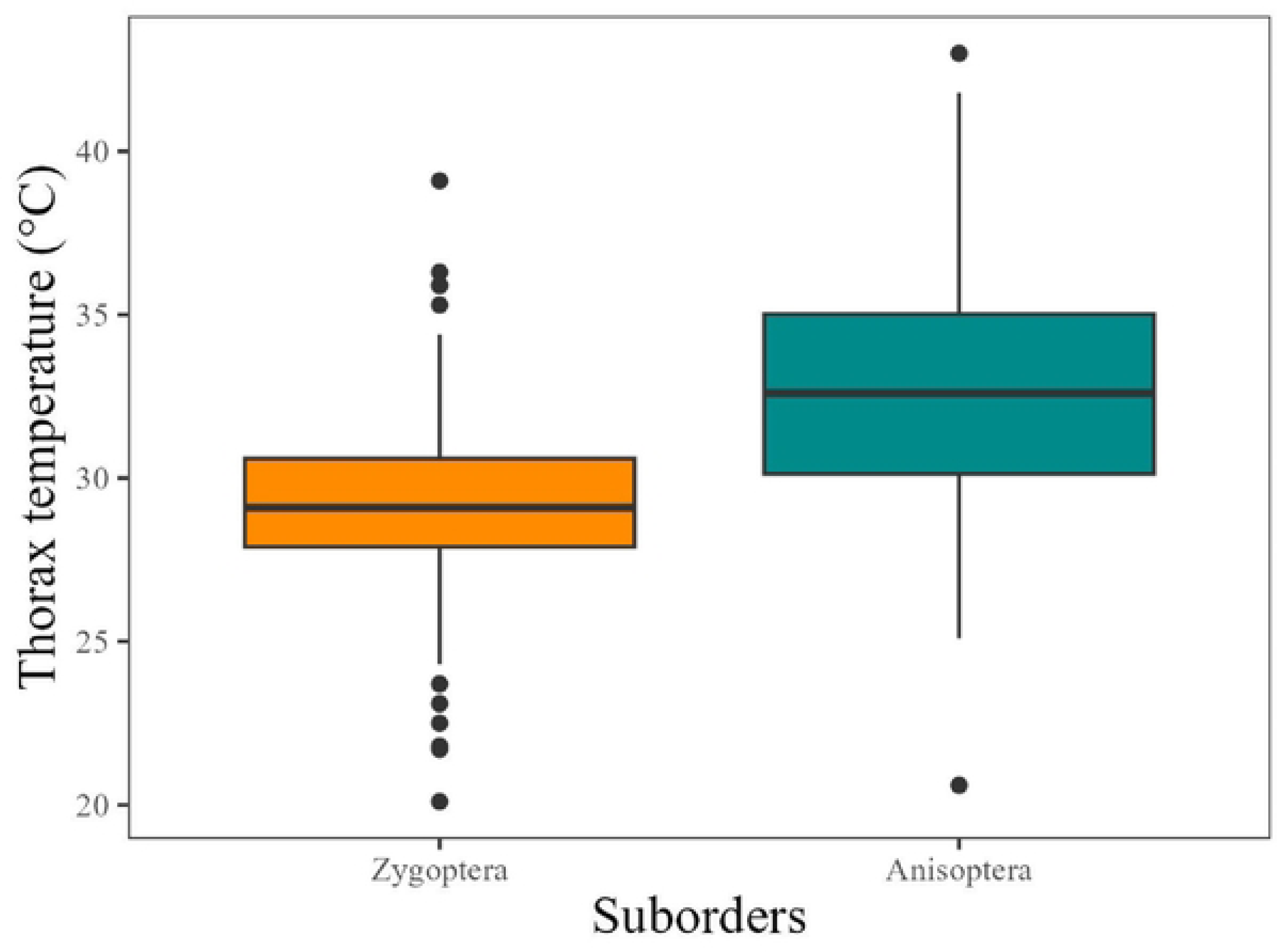
Thoracic temperature of Anisoptera (Dark orange) and Zygoptera (Cyan). Black dots are outliers.

**Table 1.** Mean and standard deviation (SD) values of thorax temperature, air temperature, and thorax width for Zygoptera and Anisoptera sampled in Brazilian Amazon streams.

| Anisoptera |  |  |  |
| --- | --- | --- | --- |
|  | Thorax width (cm) | Thorax temperature (°C) | Air temperature (°C) |
| Mean | 3.439 | 33.867 | 31.636 |
| SD | 1.465 | 3.677 | 2.013 |
| Zygoptera |  |  |  |
| Mean | 1.663 | 29.510 | 30.113 |
| SD | 0.446 | 1.603 | 1.327 |

**Table 2.** Results of the GLMM (Gaussian distribution) (species random effect) evaluating the relationship between thorax temperature of Odonata (response variable) and suborders and interaction of air temperature and thorax width.

|  | Value | Std.Error | DF | t-value | p-value |
| --- | --- | --- | --- | --- | --- |
| Intercept | 29.800 | 0.257 | 521 | 116.042 | **<0.001** |
| Suborders | 1.761 | 0.501 | 54 | 3.517 | **<0.001** |
| Thorax width (cm) | 0.820 | 0.171 | 521 | 4.809 | **<0.001** |
| Air temperature (°C) | 1.159 | 0.096 | 521 | 12.020 | **<0.001** |
| Thorax width *Air temperature | 0.288 | 0.086 | 521 | 3.361 | **<0.001** |

Significant p values appear in bold.

Anisopterans presented an average difference of air and thoracic temperature up to 8°C above air temperature and 6°C below air temperature, showing a negative relationship with thorax width (Table 3 and Fig 4). For Zygoptera, the same pattern was observed with an average difference of 3°C above and 4°C below air temperature (Table 3 and Fig 4).

**Fig 4.**
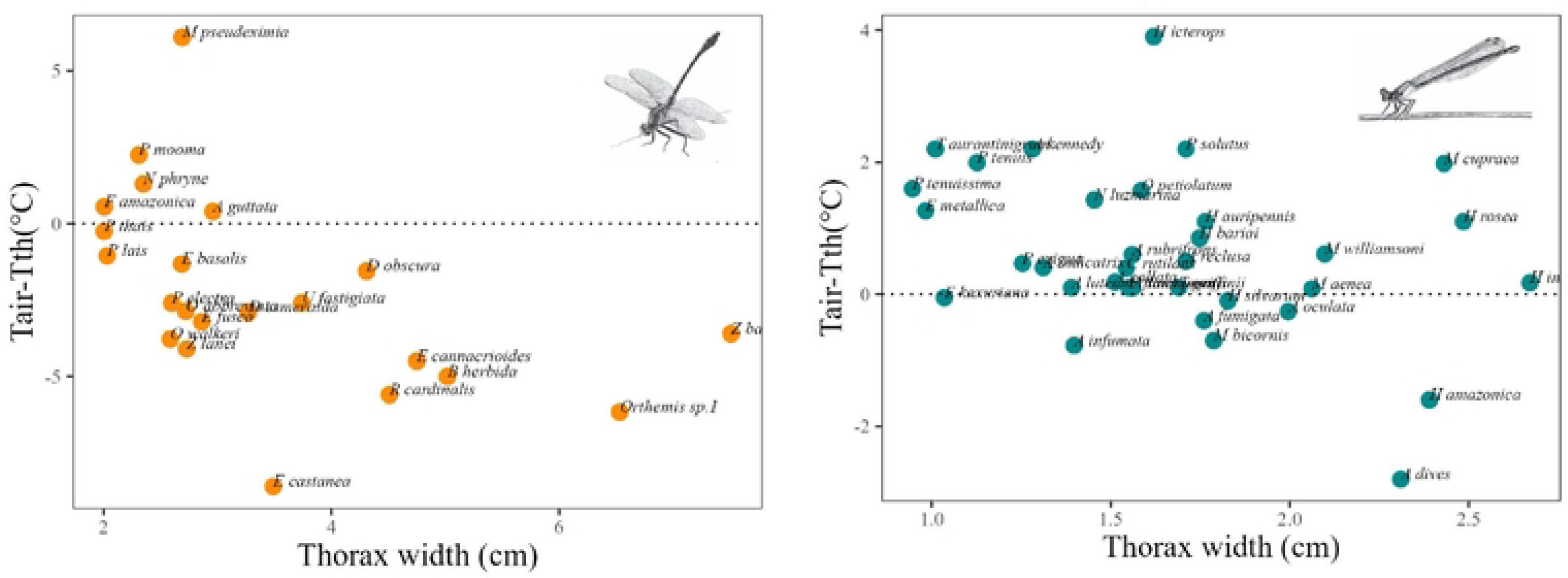
Relation of the difference between air and thorax temperature (Tair-Tth (°C) response variable) and thorax width (cm), for Anisoptera (Dark orange) dots bellow dotted line are species that have the thorax temperature above air temperature. Zygoptera (Cyan). Images of Odonata from Stehr, F.W and Tennessen, K.J.

**Table 3.** GLMM (Gaussian distribution) (species random effect) results evaluating the difference between air and thorax temperature (Tair-Tth – variable response) with thorax width of Anisoptera and Zygoptera (predictor).

| **Anisoptera** |  |  |  |  |  |
| --- | --- | --- | --- | --- | --- |
|  | Value | Std.Error | DF | t-value | p-value |
| Intercept | 0.753 | 1.034 | 192 | 0.728 | **0.468** |
| Thorax width (cm) | -0.908 | 0.290 | 192 | -3.132 | **0.002** |
| **Zygoptera** |  |  |  |  |  |
|  | Value | Std.Error | DF | t-value | p-value |
| Intercept | 2.237 | 0.578 | 330 | 3.867 | **0.000** |
| Thorax width (cm) | -0.983 | 0.341 | 330 | -2.886 | **0.004** |

Significant p value appears in bold.

### Stream structure and odonate assemblage

There was a relation between environmental variables (depth, channel width and HII) and Odonata species (p = 0.010) (OMI analysis global test). The variable that contributed the most to the first sorting axis was HII (0.88), followed by width (-0.22) and depth (0.03). This variable is essential to assess environmental integrity and demonstrates that Odonata species change composition with more intact streams (HII above 0.7) and with multiple anthropic activities (Fig 5). Streams with greater habitat integrity have a greater number of species of Anisoptera and Zygoptera that are only collected in these environments (S3 Table).

**Fig 5.**
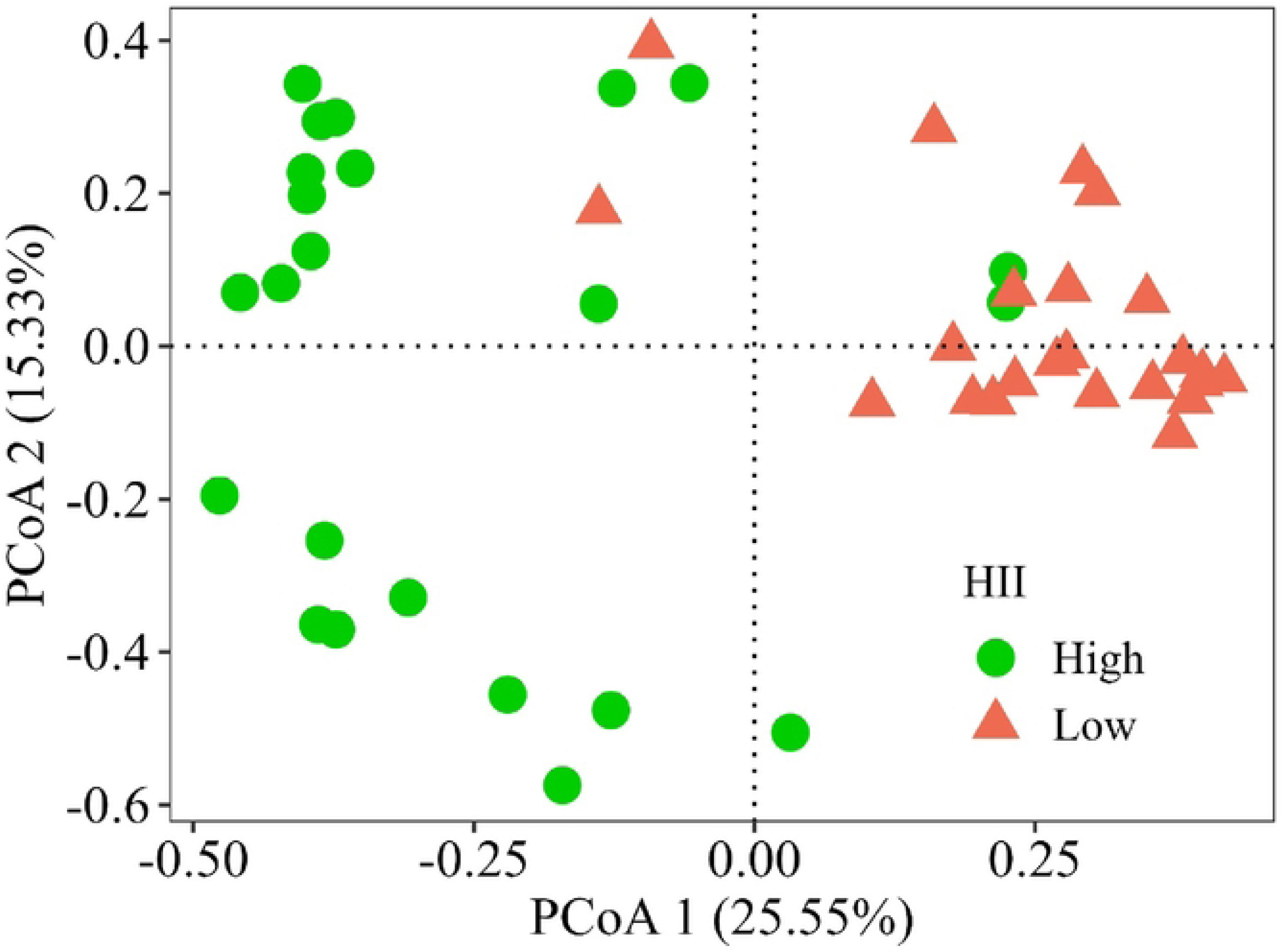
Ordination (PCoA) of Odonata species and habitat integrity index (HII): green dots are HII above 0.7 and Coral dots are streams with multiple anthropic activities (HII bellow 0.7).

### Niche and abundance of Odonata

The most abundant species of Zygoptera were *C. rutilans, M. aenea* and *E. metallica*. For Anisoptera were *F. amazonica, E. basalis* and *P. lais*. The average the niche breadth was 0.221 (standard deviation ± 0.289) and the position 2.219 (±1.94). Species with greater niche breadth were *M. cupraea, H*. *silvarum* and *M. aenea* (Zygoptera) and *E. fusca, O. walkeri* and *O abbreviata* (Anisoptera). For niche positions were *H. auripennis, A. luteum* and *H. icterops* (Zygoptera) and *D. obscura, B. herbida* and *E. cannacrioides* (Anisoptera).

Niche breadth and position were the most important predictors of Odonata abundance and suborders (Table 4 and Fig 6). Except niche position for Anisoptera. Only when evaluating suborders separately, thorax temperature emerges as important predictor for them.

**Fig 6.**
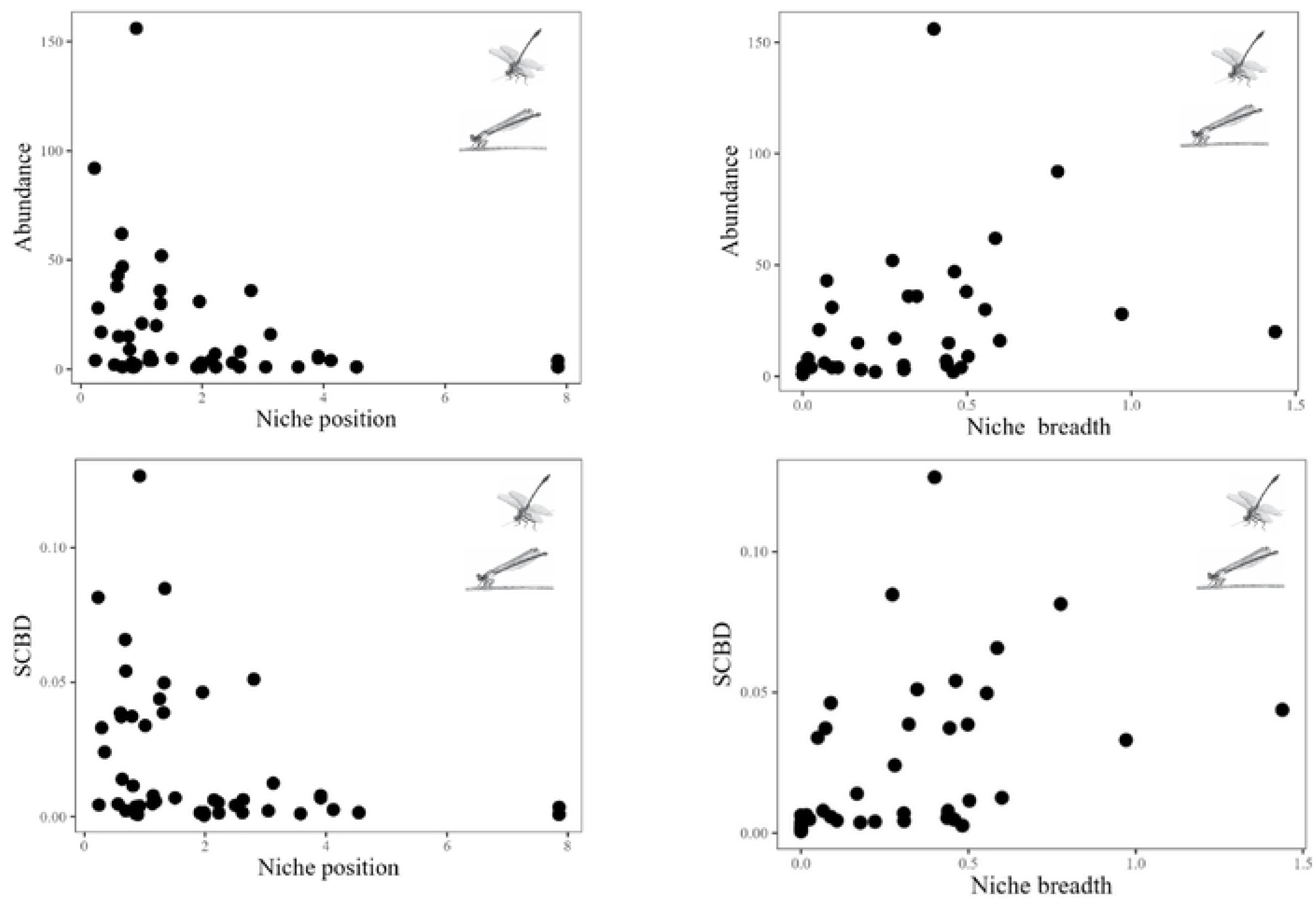
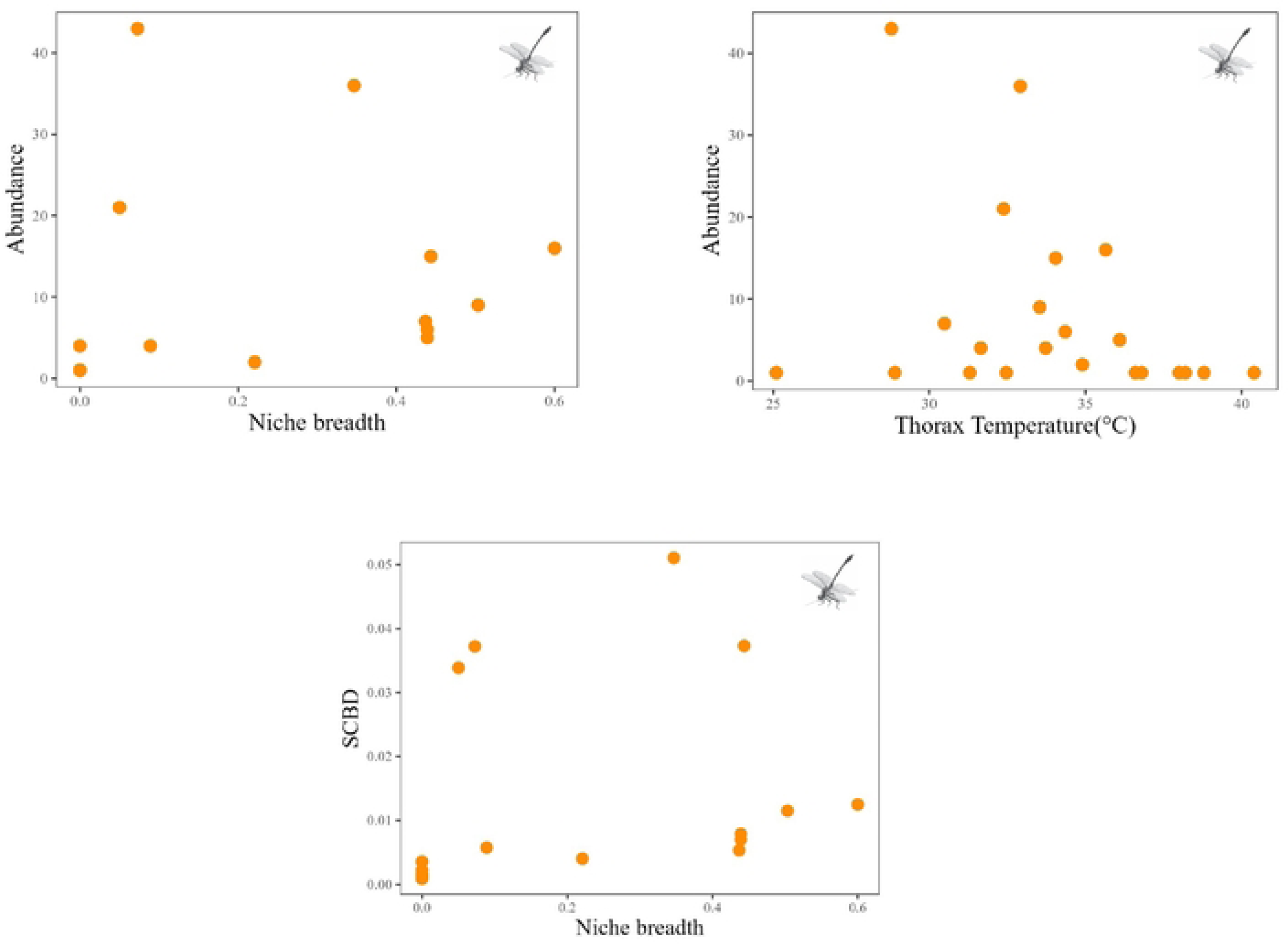
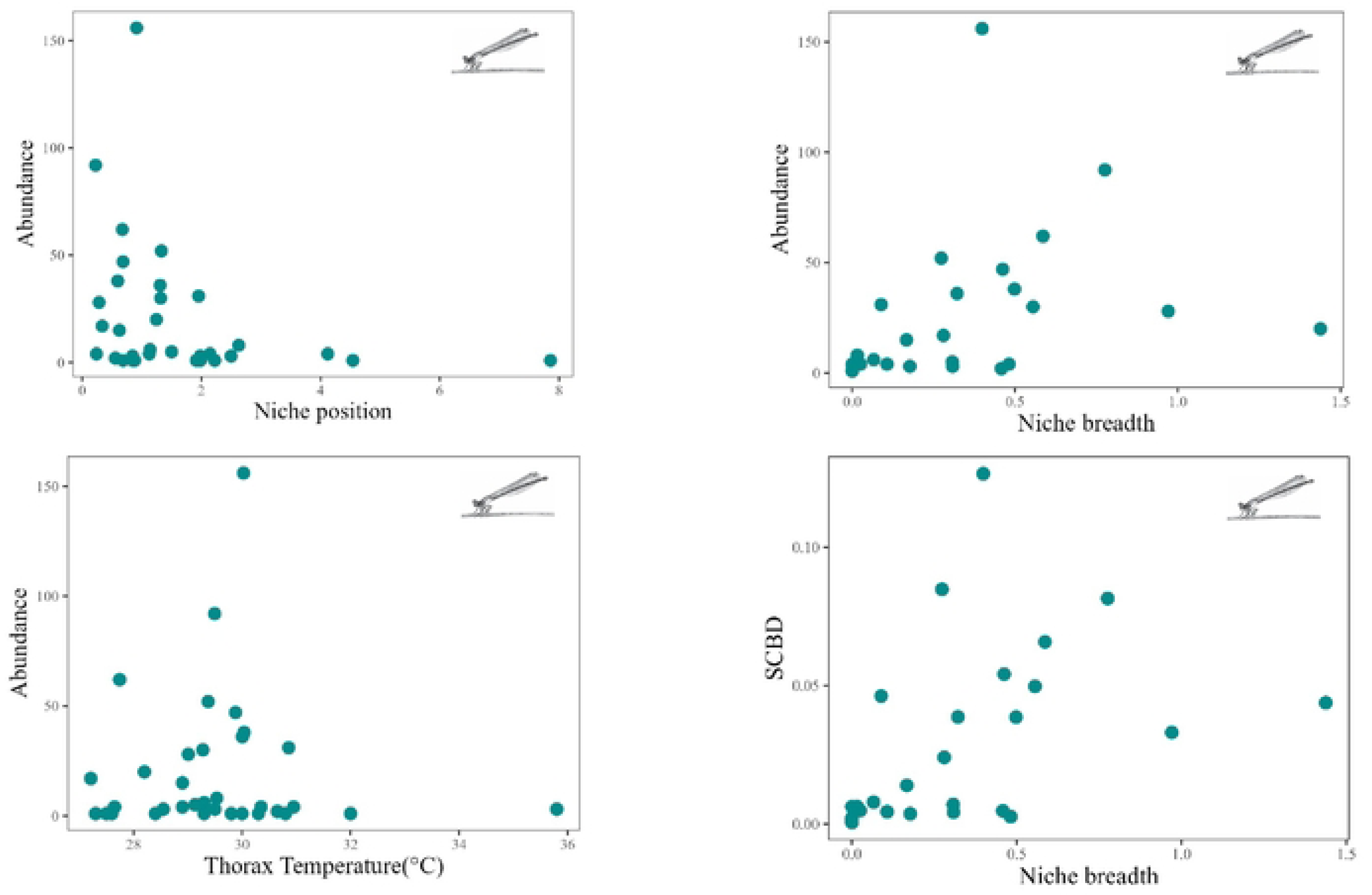
Significant relationships of Abundance and SCBD of Odonata and the suborders and niche breadth, position and thorax temperature. Odonata (Black dots), Anisoptera (Dark orange) and Zygoptera (Cyan). Images of Odonata from Stehr, F.W.

**Table 4.** GLMM results (negative binomial distribution) for Odonata (Suborder as a random effect), and GLM for suborders evaluating the relationship of species abundance with niche breadth, position and thorax temperature.

| Odonata |  |  |  |  |  |
| --- | --- | --- | --- | --- | --- |
|  | Value | Std.Error | DF | t-value | p-value |
| Intercept | 2.232 | 0.188 | 51 | 11.878 | **<0.001** |
| Niche position (NP) | -0.503 | 0.241 | 51 | -2.085 | **0.042** |
| Niche breadth (NB) | 0.856 | 0.189 | 51 | 4.529 | **<0.001** |
| Thorax temperature (°C) | -0.201 | 0.213 | 51 | -0.942 | 0.351 |
| Anisoptera |  |  |  |  |  |
|  | Estimate | Std. Error | Z Value | Pr(>\|z\|) |  |
| Intercept | 2.434 | 0.257 | 9.482 | **<0.001** |  |
| Niche position (NP) | -0.257 | 0.231 | -1.110 | 0.267 |  |
| Niche breadth (NB) | 0.850 | 0.302 | 2.817 | **0.005** |  |
| Thorax temperature (°C) | -0.612 | 0.219 | -2.798 | **0.005** |  |
| Zygoptera |  |  |  |  |  |
|  | Estimate | Std. Error | Z Value | Pr(>\|z\|) |  |
| Intercept | 2.569 | 0.285 | 9.006 | **<0.001** |  |
| Niche position (NP) | -0.886 | 0.310 | -2.854 | **0.004** |  |
| Niche breadth (NB) | 0.866 | 0.175 | 4.963 | **<0.001** |  |
| Thorax temperature (°C) | 0.868 | 0.421 | 2.064 | **0.039** |  |

Significant p values appear in bold.

On average, SCBD was 0.017 (standard deviation ± 0.026). Species with greater SCBD were *C. rutilans, H. indeprensa* and *M. aenea* for the suborder Zygoptera and *E. basalis, O. abbreviata* and *F. amazonia* for the suborder Anisoptera. When evaluating predictor variables for Odonata’s SCBD, niche breadth and position emerged as important predictor. When evaluating predictor variables for suborders SCBD, niche breadth only emerged as predictor. (Table 5 and Fig 6).

**Table 5.**
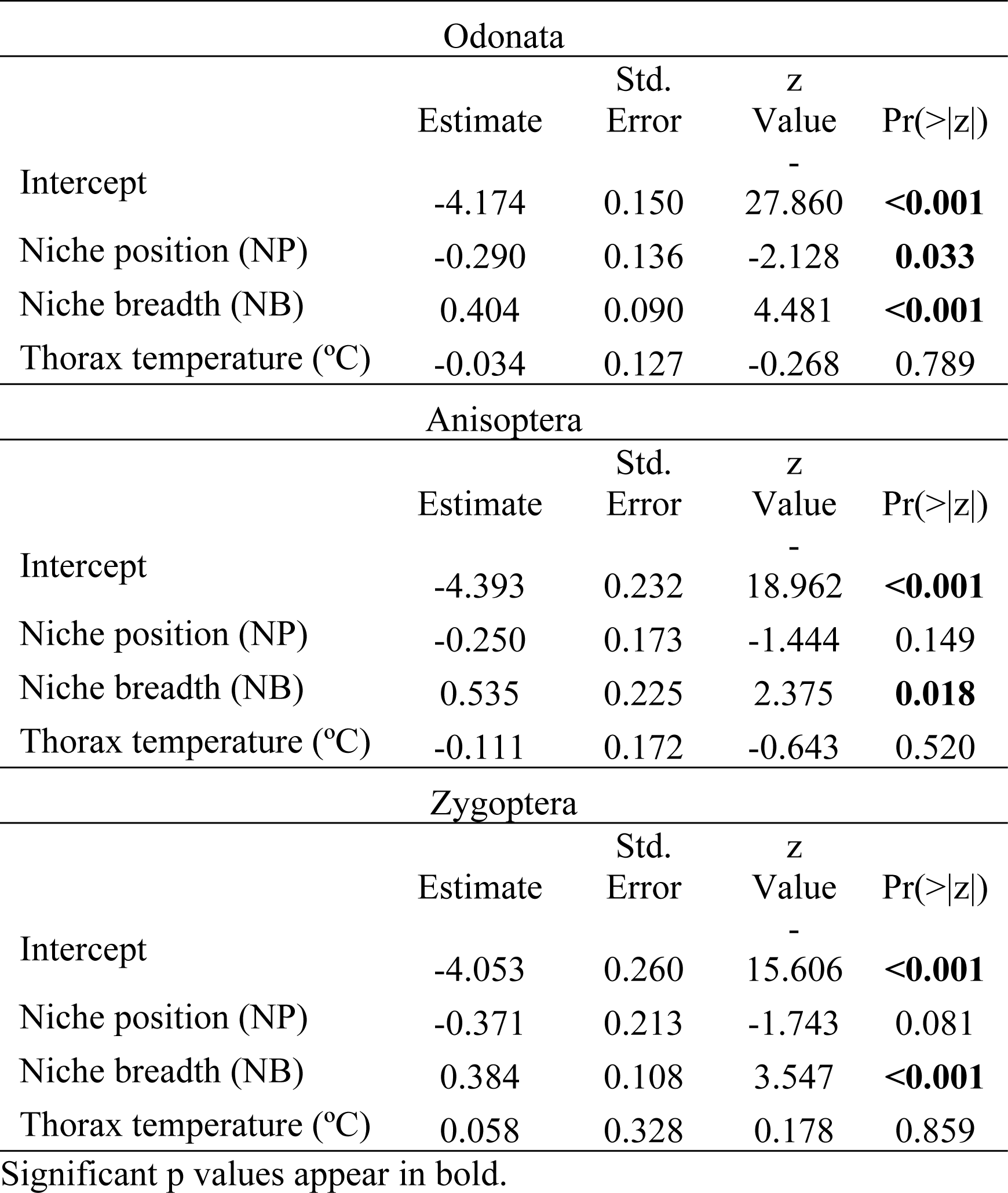
Beta regression results between SCBD of Odonata (suborders shown separately) and niche breadth, position, and thorax temperature.

## Discussion

Niche breadth was an important predictor for abundance and SCBD for Odonata and its suborders. Niche position also is important for Odonata abundance and SCBD, and Zygoptera abundance. Contrary our expected hypothesis, the thoracic temperature was a good predictor only for abundance and when evaluated suborders separately. Research shows that species with greater niche breadth are more abundant and tend to present greater regional occupation [6]. Furthermore, these species, often display a more generalist or tolerant behavior towards diverse environmental conditions, which may contribute to their reduced vulnerability. [50]. In Odonata, this macroecological pattern may be due to the generalist species being able to persist in degraded environments [51], resulting in a broader distribution range [52]. In our study the families Libellulidae and Coenagrionidae, are more abundant and both are diverse families at a continental scale which, which may explain their wide distribution [53–54]. Some species of Coenagrionidae are associated to degraded environments, within this family *E. metallica* (associated with streams with environmental change Faria et al. 2021) and *N. luzmarina*s uch as *E. fusca,* and *O. walkeri* [55–56]. In addition, other families, *M. cupraea, H. silvarum* and *M. aenea* also presented greater niche breadth. Species with lower abundance and smaller niche breadth tend to be sensitive to environmental changes, which make them less likely to persist when the physical characteristics of streams change [29].

The analysis shows that Odonata and the suborders separately, species with greater niche breadth had a greater contribution to beta diversity, which is opposite to expected [6,50]. Possibly more habitat generalist species and adults that are active dispersers and can survive in a wide range of environmental conditions play an important role in governing SCBD. Previous study show that the pattern of beta diversity of an assemblage is also adequately described by the common species, according to [58]. Only for Odonata and Zygoptera we found niche position negatively associated with abundance e and com Odonata SCBD, indicating that species, when very abundant, occur in non-marginal environments and occurs in more common habitats in the region. However, we highlight here that species such as *C. rutilans* and *M. aenea* (both contribute to beta diversity and have high niche breadth) in Amazonian streams seem to have limits to occur in different intensities of multiple impacts, as these species can disappear with impacts that lead to drastic changes in the streams (Faria et al. 2021). Contrary to our hypothesis, niche characteristics appear more important for contribution of these species to beta diversity than body temperature. On the other hand, for species abundance, thorax temperature is an important predictor, only when evaluated suborders separately. In fact, much previous work has demonstrated how environmental filters affect Odonata as a whole as microhabitat structure (Bank angle, woo in the stream bed), physicals and chemical, and canopy cover dossel (Brito et al., 2021) and regional variables surroundings streams (Luke et al. 2017).

Ecofisiological traits as body size and thorax temperature, therefore, provides additional insights into the patterns of suborders abundance and contribute to metacommunity dynamics due to its thermoregulatory restrictions [18]. There is negative relationship between thoracic temperature and abundance. In general, Anisoptera can heat their bodies through the heliothermic or even endothermic ability. These thermoregulatory abilities are crucial for Odonata distribution [18]. Furthermore, heliothermic anisopterans can benefit from habitats with reduced riparian vegetation and greater sunlight input and can be quite abundant in these areas [21]. Conversely, larger species of Anisoptera (e.g., Gomphidae) with higher body temperature may be endothermic. These species live inside forests and are often found in streams only when they arrive for mating and oviposition [59]. These ecological and behavioral traits, make this species with higher thoracic temperatures are more difficult to collect, at the same time more sensitive to loss of riparian vegetation. Zygoptera with higher temperatures are large individuals such as *H. auripennis* and *A. dives* and had lower abundance.

Thoracic temperature differs between suborders and is related to thorax width and air temperature. Anisopterans presented on average thoracic temperature up to 8°C above air temperature and 6°C bellow, showing a negative relationship with thorax width. For Zygoptera, the thorax temperature is maintained 3°C above and 4°C below air temperature. Differences in the behavior of these groups may help explain this pattern. Anisoptera tend to be heliothermic or endothermic fliers and maintain the thoracic temperature above that of the air. One of the ways to control heat loss is to alter the circulation of hemolymph between the thorax and abdomen [15], posture adjustment (Corbet, 1999). Zygoptera can maintain the temperature of the thorax closer to the air, previous studies demonstrate that this difference can be up to 1°C (Shelly, 1982). They can be conformers or in some cases and the largest being heliothermic, or even a continuum that may exist between these groups (Corbet and May, 2018) and future studies can better investigate these categories.

Finally, niche characteristics may be important for the distribution of Odonata. ecophysiological traits also was important for Anisoptera and Zygpoptera abundance. May [15] suggests also that climate, body size and behavior are essential for maintaining the body temperature of Odonata. Changes in streams due of anthropic activities alter microclimatic patterns such as air temperature, fundamental for the physiological processes of Odonata species, leading to a change in the composition of species in these environments [38]. We have demonstrated that adult odonate species composition varies in relation to habitat integrity. We therefore suggest that their monitoring would provide a good indicator of riparian zone quality considering niche characteristics and their thermorregulation habilities.

## Acknowledgments

We thank Ana Luisa Fares, Ana Luiza Andrade, Erlane José Cunha, Fernando Geraldo de Carvalho, for helping us with the biological sampling.

